# Capturing heterogeneous infectiousness in transmission dynamic models of tuberculosis: a compartmental modelling approach

**DOI:** 10.1101/2020.06.26.173104

**Authors:** Yayehirad A Melsew, Romain Ragonnet, Allen C Cheng, Emma S McBryde, James M Trauer

**Affiliations:** Department of Epidemiology and Preventive Medicine, School of Public Health and Preventive Medicine, Monash University, 553 St Kilda Road, Melbourne, VIC 3004, Australia; Department of Epidemiology and Biostatistics, Institute of Public Health, University of Gondar, Gondar, Ethiopia; Australian Institute of Tropical Health and Medicine, James Cook University, Townsville, QLD 4811, AUSTRALIA

**Keywords:** heterogeneous infectiousness, M. tuberculosis transmission, mathematical model

## Abstract

Infectiousness heterogeneity among individuals with tuberculosis (TB) is substantial and is likely to have a significant impact on the long-term dynamics of TB and the effectiveness of interventions. However, there is a gap in capturing heterogeneous infectiousness and evaluating its impact on the effectiveness of interventions.

Informed by observed distribution of secondary infections, we constructed a deterministic model of TB transmission using ordinary differential equations. The model incorporated assumption of heterogeneous infectiousness with three levels of infectivity, namely non-spreaders, low-spreaders and super-spreaders. We evaluated the effectiveness of dynamic transmission untargeted and targeted implementation of an intervention intended to represent active case finding with a point-of-care diagnostic tool. The simulated intervention detected 20% of all TB patients who would otherwise have been missed by the health system during their disease episode and was compared across four epidemiological scenarios.

Our model suggested that targeting the active case finding intervention towards super-spreaders was more effective than untargeted intervention in all setting scenarios, with more effectiveness in settings with low case detection and high transmission intensity. For instance, a targeted intervention achieved a 42.2% reduction in TB incidence, while the untargeted intervention achieved only a 20.7% reduction over 20 years, given the same number of people treated. Although the most marked impact on equilibrium TB incidence came from the rate of late reactivation, the proportion of super-spreaders and their relative infectiousness had shown substantial impact.

Targeting active case-finding interventions to highly infectious cases likely to be particularly beneficial in settings where case detection is poor. Heterogeneity-related parameters had an equivalent effect to several other parameters that have been established as being very important to TB transmission dynamics.

## Introduction

Tuberculosis (TB) is the world’s leading cause of death from a single infectious agent, in 2019, ranking above HIV/AIDS with an estimated 10.0 million cases and 1.2 million deaths worldwide [1]. The World Health Organization developed the new End TB Strategy for post-2015 TB elimination activities, with the ambition of a 95% reduction in TB deaths and a 90% reduction in TB incidence by 2035 by comparison to 2015 rates [2]. However, the natural history of TB remains poorly understood, and consequently, the uncertain potential impact of control interventions limits confidence about the possibility of its elimination. The heterogeneous transmission of *Mycobacterium tuberculosis (Mtb)* within populations is well-established, but its epidemiological impact is poorly understood. Moreover, up to a third of TB cases are not diagnosed, so finding and treating infectious cases is key to achieving TB control, and finding and treating highly infectious people is the key to TB control in high transmission settings [3, 4].

We define infectiousness heterogeneity as the variability in the capacity of infectious patients to produce secondary infections, which may be attributable to characteristics of the host, the agent and the environment [5–9]. Heterogeneity of infectious individuals in spreading *Mtb* infection is well-recognised, with a small group of highly infectious individuals producing a large proportion of secondary infections, while many others produce very few or none [10]. Previous TB genomic epidemiology has identified TB super-spreading events by quantifying heterogeneity in the infectiousness of TB patients through fitting a standard statistical distribution (the negative binomial distribution, NBD) to the distribution of secondary cases produced by each infectious patient (the “offspring distribution”) [11]. More recently, data from TB contacts in a low-transmission setting have allowed estimation of the proportion of all TB patients that can be categorised as super-spreaders, finding it to be approximately 10% [9]. In such studies, super-spreaders were typically defined as patients who produced a number of secondary “infections/cases” greater than the 99^th^ centile of a standard Poisson offspring distribution, with distribution mean equal to the average number of secondary infections per index [9, 12]. This heterogeneity is likely to have implications for both the burden of disease and the effectiveness of control interventions. Thus, we propose that it is necessary to capture this heterogeneity when modelling TB transmission dynamics.

Some past compartmental models of TB transmission dynamics have attempted to capture heterogeneity in patients’ infectiousness by stratifying the active TB compartment into different levels of infectivity. Amongst the most typical approaches is stratification as either infectious (usually representing pulmonary TB) or non-infectious (extrapulmonary TB) [13–17]. This approach implies that all pulmonary TB patients are equally infectious, while all extrapulmonary patients are entirely non-infectious, and so does not capture the heterogeneity among pulmonary patients. Other TB models have considered sputum smear status as a factor in stratifying patients’ levels of infectiousness, considering both smear-negative and smear-positive pulmonary TB to be infectious, with the relative infectiousness of smear-negative patients compared to smear-positive typically set between 15 and 25% [18–33]. While stratifying pulmonary TB patients based on smear status captures an additional clinical attribute that is important in determining infectiousness, even these models still do not capture the full picture of TB patients’ infectiousness variation, since several behavioural and demographic factors other than smear status can affect the level of infectiousness [5].

An individual-based model simulated patterns of meetings and *Mtb* transmission between three different types of contact to determine the effects of variation in infectiousness and susceptibility on transmission location. The study defined super-spreading defined based on the number of secondary cases instead of secondary infections and suggested that the majority of disease resulted from infection by a small proportion of people with TB or super-spreaders [34]. However, as TB disease activation may take long time and also depend on mainly on the characteristics of contact person, the definition of super-spreading used in the study may not show true heterogeneity in *Mtb* transmission [35]. In the current study, we present a deterministic compartmental *Mtb* transmission model that incorporates three levels of TB patients’ infectivity, namely non-spreaders, low-spreaders and super-spreaders. Using this framework, we analysed how infectiousness heterogeneity affects disease dynamics and evaluated the effectiveness of a hypothetical active case finding intervention and the impact of targeting super-spreaders.

## Methods

### Literature review

Before constructing our model, we systematically reviewed how previous TB models have approached heterogeneous infectiousness [36]. In the review, we found that TB models frequently stratified the active TB compartment according to one or more patient-related factors. We constructed a flexible model with three levels of infectiousness, without restricting to a given factor, and the ability to transition between these levels using empirical measures of heterogeneous infectiousness to parametrise the model.

### Empiric data

Data from the VTP which are stored by the Victorian Department of Health and Human Services (DHHS) were used for the following analysis. Index patients were classified as confirmed cases of TB notified from 1 January 2005 to 31 December 2015 in residents of Victoria. The data set includes contact tracing information and results of testing for *Mtb* infection, with cases of subsequent active TB disease linked to these contact episodes now extending to March 2017 (see [37] for earlier publication of linkage process). We constructed empirical offspring distributions from the detailed contact tracing data set of the VTP.

Ethical approval was obtained from Monash University, Human Research Ethics Committee (Project Number: 7776) and permission was given by the VTP and DHHS.

### Model structure

Using ordinary differential equations (ODE), we constructed a deterministic model of *Mtb* transmission in a hypothetical high TB burden setting. To represent heterogeneous infectiousness and super-spreading in *Mtb* transmission, we stratified the active TB compartment into three sub-classes with different levels of infectiousness: non-spreaders (*I* _0_), low-spreaders (*I* _1_) and super-spreaders (*I* _2_). We did not use any specific factor to discriminate *I* _0_, *I* _1_ and *I* _2_ a priori. Instead, we use the offspring data (number of secondary infections per index) and a Bayesian approach to find the most realistic parameterisation to implement heterogeneous infectiousness in our model.

**Fig 1** presents the model structure. The model simulates a closed population, such that births (*π*) accrue into the fully susceptible compartment (*S*) to replace all deaths. Natural death occurs from all compartments at a constant rate (*μ*). Susceptible individuals are dynamically infected, with the force of infection (*λ*) contributed by super-spreaders and low-spreaders. Following infection, individuals enter a rapid-sojourn early latent compartment (*L*_*A*_), from which they may progress to one of the three active TB compartments (*I* _0_, *I* _1_ or *I* _2_) at a total rate of *ε* or enter the low-risk late latent stage (*L*_*B*_) at rate *κ*. From the late latent state they may progress more slowly to disease (total rate *ν*), also entering the active TB compartments ¿). Re-infection may occur during late latency with a reduced force of infection *λ*_*r*_ (*λ*_*r*_=*r × λ*) and, if re-infection occurs, these individuals similarly enter the early latent compartment, progressing to the disease at the same rates as for newly infected persons. Individuals with active disease may either die due to TB (rate *μ*_*i*_) or background mortality (*μ*); or spontaneously recover and return to the late latent state (rate *γ*); or be detected and treated by the health system (rate *δ*) and return to the susceptible compartment (*S*).

**Fig 1:**
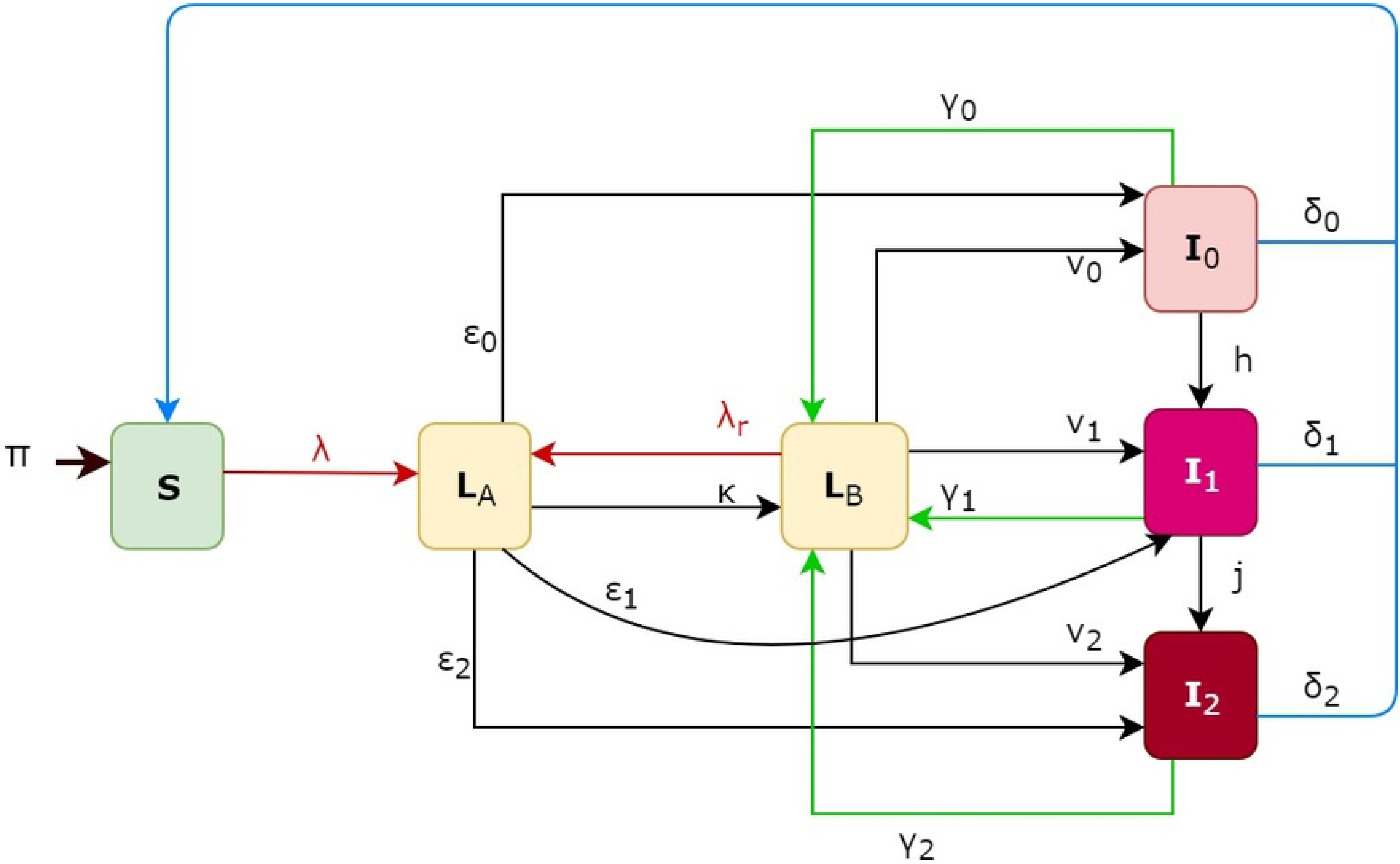
Transmission dynamic model of TB with heterogeneous infectiousness. Boxes represent compartments in the model and arrows represent flows. Not shown here but included in the model are natural mortality from each compartment and TB-related mortality from each active TB compartment (*I* _0_, *I* _1_ and *I* _2_). Red arrows represent transmission flows; black arrows represent disease progression processes; green arrows represent self-recovery, blue arrows represent detection and treatment cure.

### Estimation of the infectiousness parameters using empiric evidence

The four model parameters that capture heterogeneous infectiousness in the model were the proportions of non-spreaders, low-spreaders and super-spreaders, and the relative infectiousness of super-spreaders compared to low-spreaders. In order to estimate these parameters, we first determined the statistical distribution of the number of secondary infections per index associated with the stochastic equivalent (continuous-time Markov) model of our ODE-based model. We obtained a mixture of three classic statistical distributions: two geometric distributions for low-spreaders and super-spreaders and one Dirac delta distribution for non-spreaders. A detailed demonstration of this result is provided in the ***Supplementary Material***.

A Hamiltonian Monte Carlo (HMC) algorithm was used to generate 20,000 samples from the posterior distributions of the parameters. The 20,000 samples used for inference were obtained from 40,000 iterations of the HMC, burning the first 20,000 draws. Burn-in size and convergence were assessed by inspecting the trace plots of the estimated parameters. The reported 95% credible intervals were obtained by computing the 2.5^th^ and 97.5^th^ percentiles of the parameters’ posterior distributions. The parameter estimates from the Bayesian process is provided in **Supplementary fig.1**.

The resulting statistical distribution was then compared in a Bayesian context with empiric data obtained from our previous analysis of TB contact tracing data in the Australian state of Victoria [9]. A comparison between the distribution of the number of secondary infections per index simulated by our calibrated model with the one observed empirically is presented in Fig 2.

**Fig 2.**
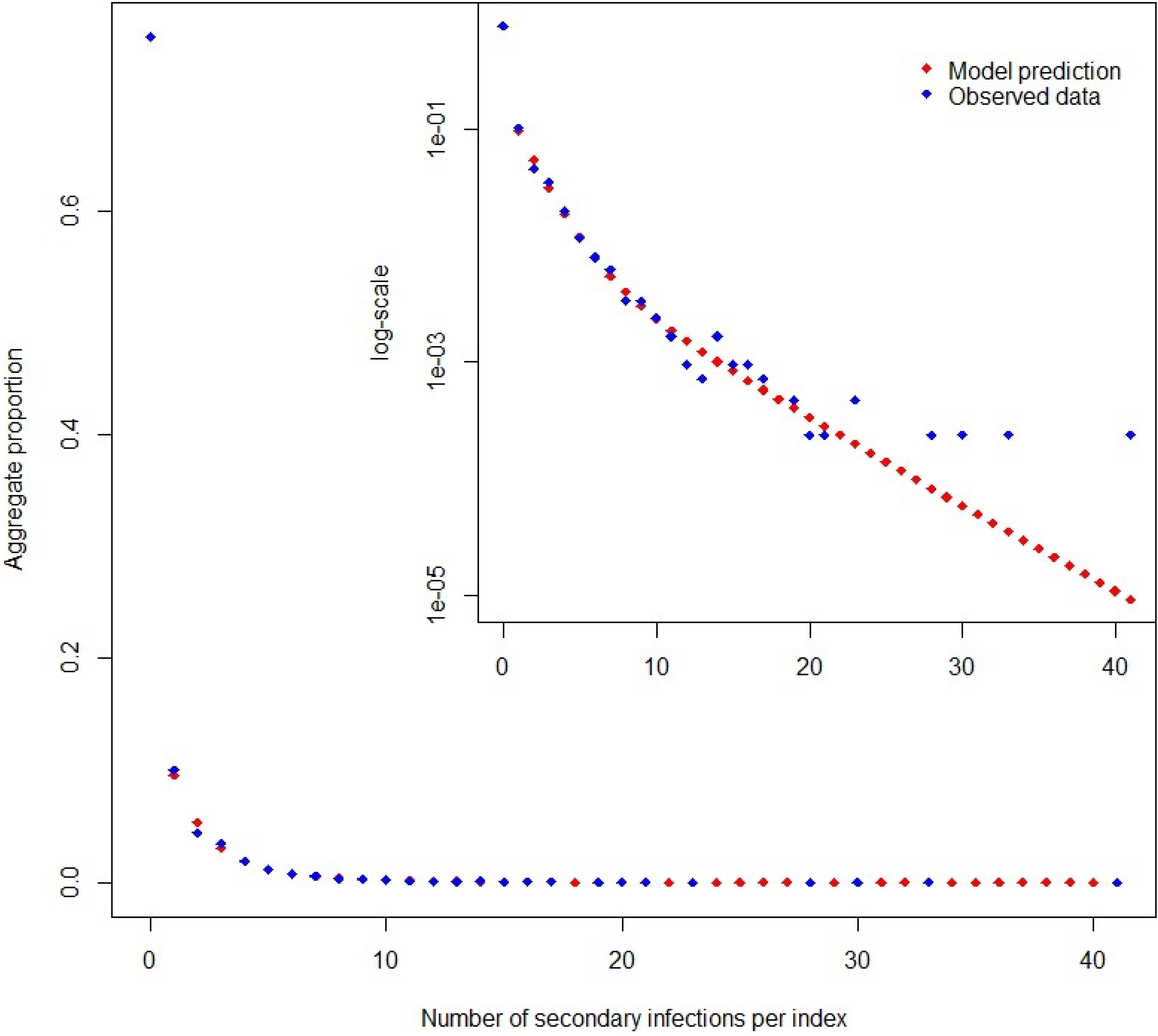
Comparison of the effective modelled secondary infection distribution with empiric data from Victoria, Australia (red points represent the modelled distribution while blue points are observed empiric data).

The parameter values other than those pertaining to infectiousness introduced above were estimated from a review of relevant evidence, with the relevant sources presented in **Table 1**.

**Table 1:**
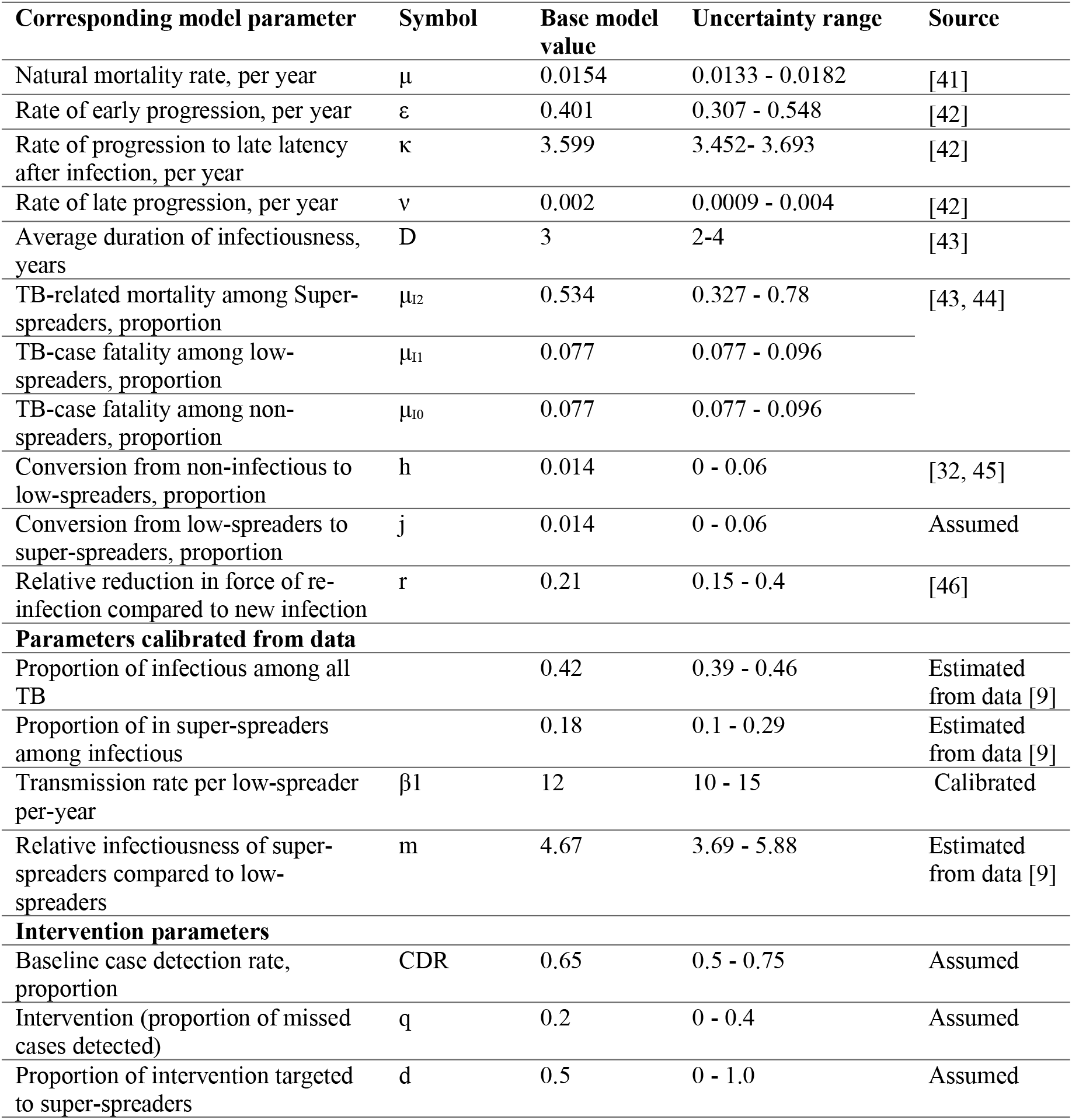
Model parameter values

### Model calibration

For the baseline equilibrium state, the ODE model was calibrated to a high TB burden settings with an incidence rate of 200 per 100,000 population per year by adjusting the transmission rate parameter for low-spreaders.

The model was programmed in R 3.4.0 (R Project for Statistical Computing), and we used a package called ‘deSolve’, solver for initial value problems of ODE [38, 39]. The Bayesian parameter estimation was done using the package ‘rstan’ (v.2.18.2) [40] and the source code for the model is publicly available from https://github.com/Yayehirad/TB_heterogeneity

### Sensitivity analysis

We first performed a one-way sensitivity analysis on each parameter to understand the sensitivity of equilibrium incidence to parameter ranges in the absence of interventions. We then undertook multidimensional sensitivity analyses to assess the correlation between equilibrium incidence and each parameter by taking 1000 samples from each parameter ranges with Latin hypercube sampling (LHS) technique [47, 48].

### Intervention simulations

We simulated an intervention intended to represent active case finding with a point-of-care diagnostic tool, accelerating successful diagnosis and treatment of individuals as early as possible before they start transmitting the infection. This could be conceptually considered as implemented by a highly sensitive point-of-care diagnostic technology following community awareness campaigns that target individuals with noticeable symptoms in order to increase their rate of presentation [49]. To target the intervention, clinical symptoms and behavioural characteristics can be used to identify those individuals who are more likely to be super-spreaders.

We implemented an intervention capable of finding *q* proportion of TB cases that would otherwise remain undetected by the passive case detection process: i.e. *CDR*_*intervention*_ =*q×*(1−*CDR*_*baseline*_)+ *CDR*_*baseline*_, where, CDR is the case detection rate, being the proportion of all cases detected during their disease episode and *q* is the proportion of cases detected under the intervention scenario that would otherwise be missed by the baseline CDR. We set the maximum attainable level of CDR in all intervention scenarios including targeting super-spreaders at 90%. In targeting the intervention towards super-spreaders, the maximum possible CDR targeted to super-spreaders under the intervention occurs when the CDR for non-spreaders and low-spreaders is kept at the baseline. With this restriction, we defined a single parameter (*d*) to represent the extent of intervention targeting relative to the maximum amount of targeting possible, which ranges from zero to one, with zero representing untargeted intervention and one representing 100% targeting. That is, the CDR targeted to super-spreaders under the intervention scenario was *CDR*_*intervention*_ plus *d* multiplied by the difference between the maximum possible CDR and *CDR*_*intervention*_. This implementation ensures equivalent effort or the number of people treated for untargeted and targeted interventions; details of these calculations are given in the ***Supplementary Material***.

We then evaluated the effectiveness of targeting super-spreaders (*d* >0) compared to untargeted intervention (*d*=0) in four independent scenarios. These four scenarios were settings with high transmission and high case detection, high transmission and low case detection, low transmission and high case detection, and low transmission and low case detection. These four scenarios were simulated by calibrating the transmission parameter for low-spreaders and CDR. In addition, we evaluated the impact of levels of targeting (*d*) for different coverage levels of the intervention (*q*) in these four setting scenarios, while other model parameters were sampled from the same plausible ranges as in the baseline sensitivity analysis using LHS.

## Results

### Baseline sensitivity analyses

The sensitivity analyses show that the most marked impact on equilibrium incidence arose from the rate of late TB progression followed by the CDR **(Fig 3).** The proportion of super-spreaders among persons with infectious TB and the relative infectiousness of super-spreaders compared to low-spreaders also had a substantial impact. **Supplementary Fig 2** presents a multidimensional sensitivity analysis, using the LHS method to sample 1000 parameter sets, which was consistent with the results of the one-way sensitivity analysis.

**Fig 3:**
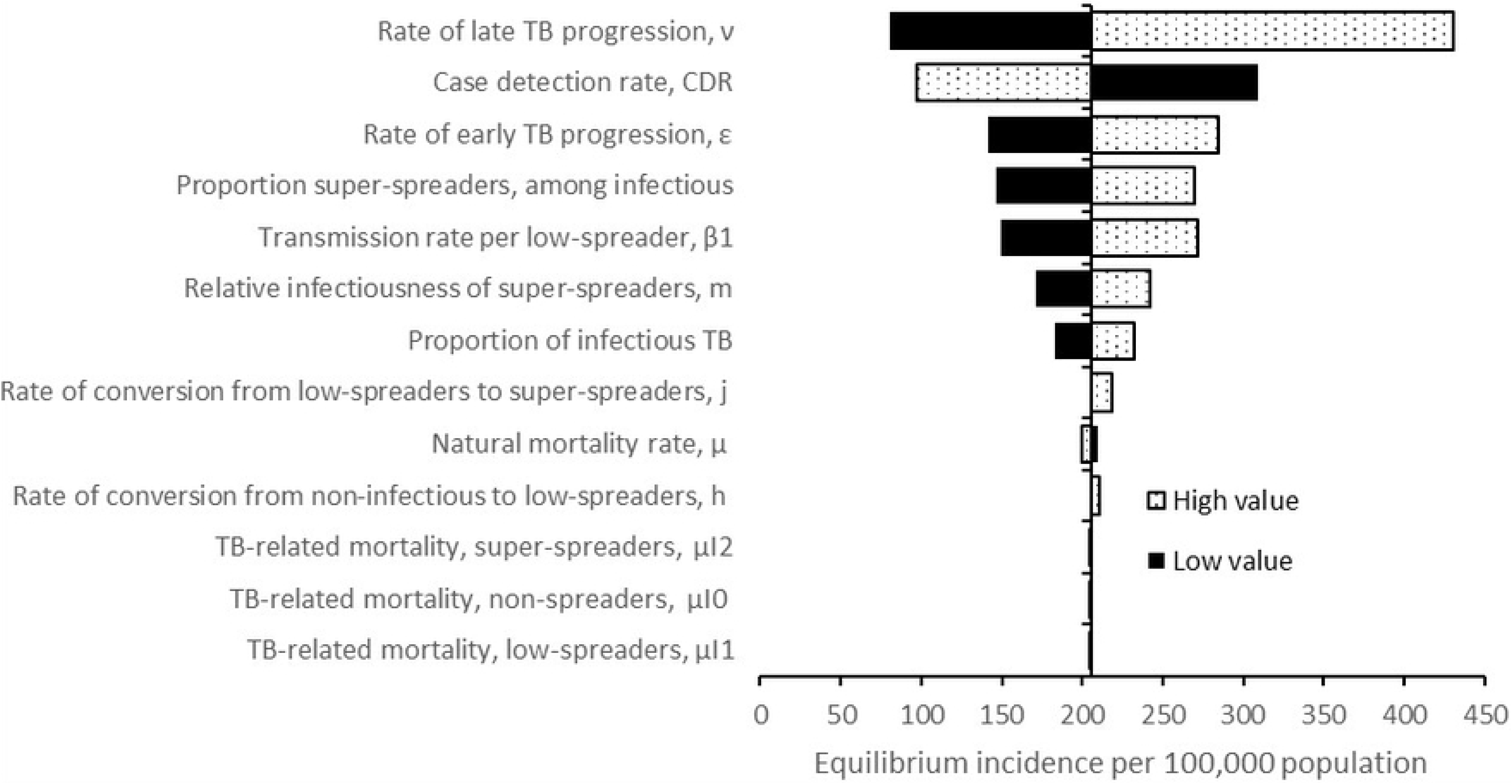
One-way sensitivity analyses of equilibrium TB incidence to variation between extremes of plausible parameter values. Maximum and minimum parameter values are given in Table 1. Values on the x-axis represent the equilibrium TB incidence, and the vertical line indicates the equilibrium incidence obtained with the baseline parameter set.

### Impact of targeted interventions

Targeting the active case finding intervention towards super-spreaders was more effective than mass intervention in all setting scenarios. However, its effectiveness was particularly marked in settings with low case detection, regardless of transmission rates. For example, in the first scenario of both high transmission and high CDR, a 20% untargeted active case finding intervention led to a 22.8% reduction in incidence over 20 years, while with an 80% targeting an equal active case finding intervention resulted in a 35.9% reduction in TB incidence over the same period. However, in the scenario of high transmission but low CDR, the targeted intervention is much more effective (achieving a 42.2% reduction) than the untargeted intervention, which only reduced incidence by 22.8% over 20 years, considering a 20% active case finding and an 80% level of targeting. Interventions in settings of low transmission and high case detection rates had a relatively minor impact on the burden (***Supplementary Fig. 3***).

We evaluated the impact of intensifying active case finding (*q*) from zero to 40% detection and varying the proportion of this intervention targeting the super-spreaders (*d*). As shown in **Fig 4**, both these quantities are essential – i.e. the increase in CDR and the extent of targeting provided the increase in CDR is more than negligible. The most significant impact of targeting was in the setting of high transmission and low CDR at all coverage levels of the intervention. Targeting super-spreaders was more effective at all levels of relative CDR increases in settings with high transmission and low case detection (Panel B), while in settings with high transmission and high case detection, targeting super-spreaders has a significant effect if the relative increase in CDR is low (Panel A).

**Fig 4:**
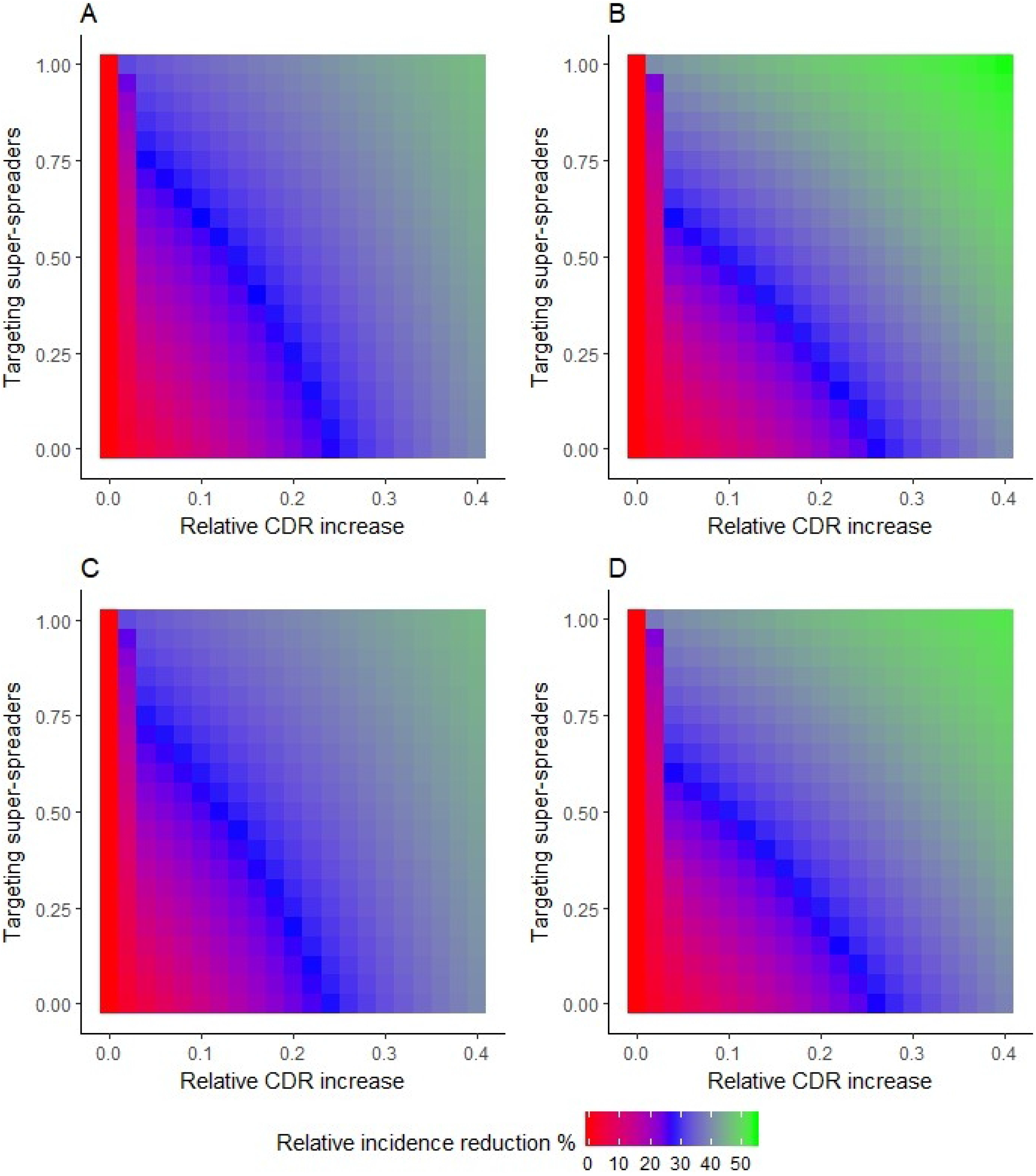
Comparison of extent of active case finding and the proportion of targeting to super-spreaders on the 20-year projected relative reduction in TB incidence under the four scenarios. A) High transmission and high case detection setting; B) high transmission and low case detection; C) low transmission and high case detection; D) low transmission and low case detection.

**Fig 5** presents the results of the one-way sensitivity analysis performed to observe the impact of high and low extremes of parameter values on the intervention’s 20-year projected relative incidence reduction. The sensitivity analyses show that the most marked impact on intervention effectiveness arose from the rate of late progression, followed by the rate of early progression and the proportion of super-spreaders. Notably, while higher rates of late progression were correlated with lower effectiveness of the active case finding intervention, higher rates of early progression resulted in greater reductions in the incidence.

**Fig 5:**
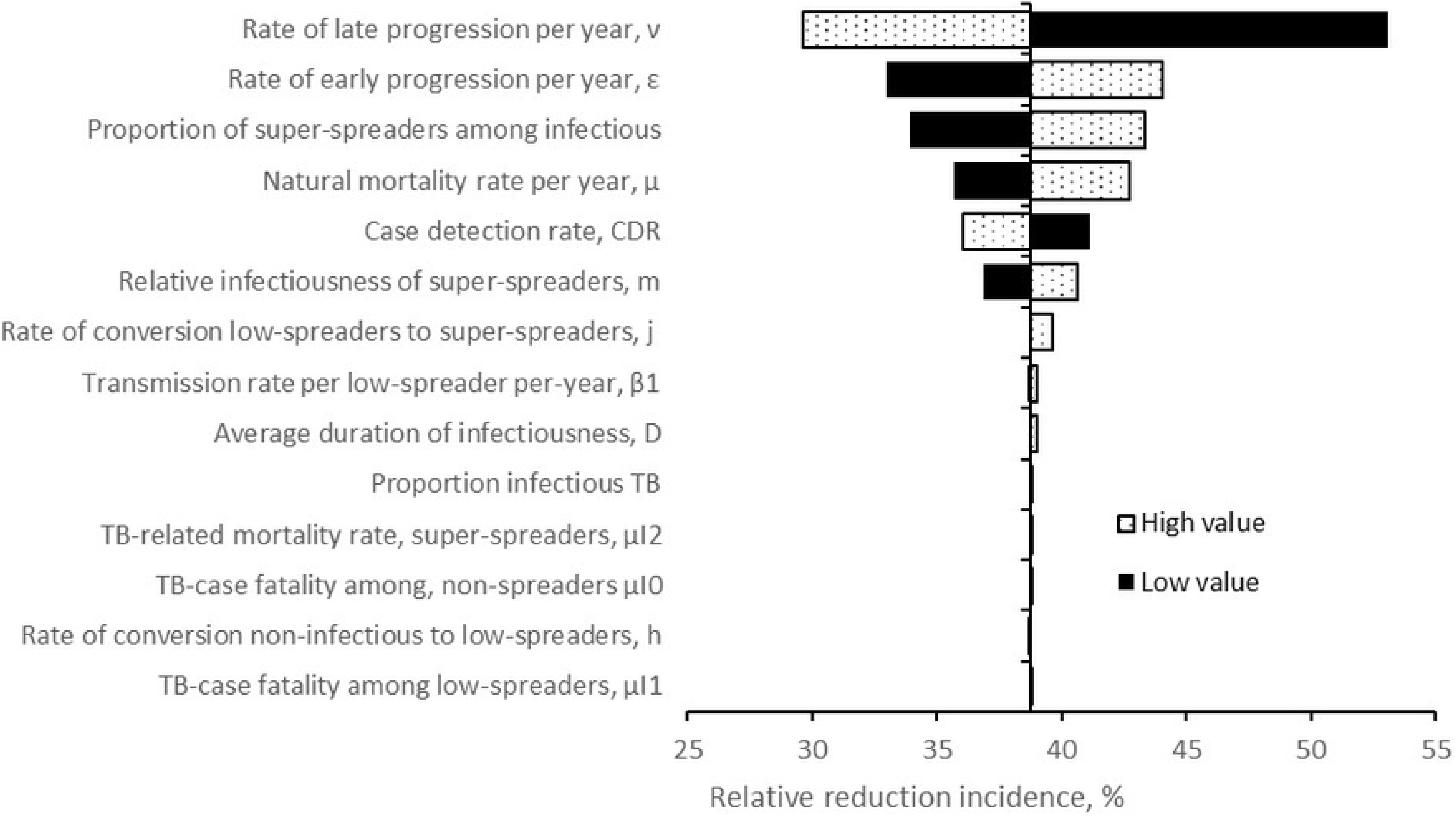
One-way sensitivity analyses of 20-year projected relative reduction in incidence to variation between extremes of plausible parameter values. The maximum and minimum parameter values are given in Table 1. Values on the x-axis represent the relative reduction in incidence following the implementation of a 20% active case finding intervention with 80% targeting. The vertical line indicates the relative reduction in incidence obtained with the baseline parameter set.

**Supplementary fig 4** presents the results of the multidimensional sensitivity analyses assessing the impact of parameter ranges on intervention effectiveness (relative reduction in annual incidence with 20-years projections). This multidimensional analysis supports the results of the one-way sensitivity analysis: that the proportion of super-spreaders had a considerable impact on intervention effectiveness, and was the most important parameter after the disease progression parameters.

## Discussion

Our model suggested that targeted active case finding interventions directed toward people likely to be super-spreaders could have a substantial impact on the TB burden, particularly in settings with high transmission but low case detection rates. This model also showed that parameters related to infectiousness heterogeneity such as the proportion of super-spreaders and their relative infectiousness compared to other low-spreaders were as crucial as other previously recognised epidemiological parameters, such as the latency progression parameters, in determining the burden of TB. The current model is also able to incorporate a range of commonly used approaches to capturing heterogeneous infectiousness of active TB cases. The choice of three infectiousness levels for active TB and the values assigned to each compartment was based on empirical data estimates, strengthening the validity of our findings.

Active case finding has been described as “turning off the tap” in TB control intervention strategies since it represents a method of identifying individuals with TB and promptly initiating treatment to avoid further onward transmissions [50]. Previous TB modelling has suggested that active case finding with highly sensitive diagnostic tools can reduce delay to treatment and so significantly reduce TB transmission [51]. The use of point-of-care diagnostic technologies with high sensitivity to detect persons with TB soon after they begin to transmit, such as GeneXpert Omni [49], could enable the implementation of active case finding interventions similar to that considered in this study. In fact, any TB active case finding intervention necessarily incorporates some level of targeting towards the highly infectious individuals, because highly infectious patients’ disease characteristics (such as higher sputum-bacilli concentration and lung-cavitation) make them more easily detectable with existing microbiological and radiological diagnostic tools. This means that implementing more sensitive diagnostics may paradoxically decrease targeting of super-spreaders – whereas traditional smear-based interventions might ensure that resources are targeted to the most infectious.. In agreement with previous modelling [52], our model suggested that active case finding interventions are particularly valuable in high transmission and low case detection settings to limit onward transmission. In the implementation of the intervention, although we did not perform cost-effectiveness analyses, our study shows that finding super-spreaders (less than 10% of all TB cases) is equally as effective epidemiologically as finding 40% of general TB population. In addition, the cost of identifying and treating super-spreaders that only comprises less than 10% of total TB cases can be compared with the cost of treating all active TB cases in a population. Our findings concerning the effectiveness of interventions can provide broad directions to future TB modellers and policymakers, although predictions from our model are not intended to provide location-specific estimates.

Our sensitivity analyses showed that a higher late reactivation rate had a negative impact on the effectiveness of active case finding interventions, while higher rates of early progression increased the impact on the 20-year projected reduction of incidence following the intervention. This suggests that active case finding interventions are more effective if the TB epidemic is more dependent on early progression and intense recent transmission than late TB reactivation since the intervention would have a more significant effect on rapidly reducing transmission. A recent study showed that variation in the latency progression parameters had important impacts on model predictions around the effectiveness of preventive therapy interventions [53], while our analysis complements this finding by showing that these parameter variations also have indispensable effects on predictions regarding the effectiveness of case-finding interventions.

In our analyses, the rate of conversion from low-spreader to super-spreader had little significance, and impact of the rate of conversion from non-infectious to low-spreaders was negligible. It is known that active TB cases’ ability to spread *Mtb* may increase as they progress clinically to more severe disease, e.g. from smear-negative to smear-positive or from non-cavitary to cavitary-TB [54]. In previous TB models, spontaneous conversion from less infectious to more infectious states such as smear-negative to smear-positive was included with rates around 1.5% per year [18, 25, 32, 45, 55–57]. These parameters do not affect model outputs substantially, such that they can be omitted for model simplicity if the actual values are consistent with what has previously been assumed. Nevertheless, the exact values of these quantities remain uncertain, and future research to refine these quantities may modify this conclusion.

The current model structure enhances flexibility around the assumption of infectiousness heterogeneity that allowed for reflection of a broad range of factors that can alter active TB cases infectivity level [5, 58]. Thus, the model is able to incorporate many previous compartmental modelling structural approaches, including those that stratified active TB cases’ infectivity into two levels, as non-infectious and infectious [13–17], or model structures that incorporate three levels of infectivity, as non-infectious, smear-negative infectious and smear-positive infectious [30, 31, 59]. However, in the application of this model to a particular epidemiological setting, data to estimate the proportion of super-spreaders is essential, since this information has a particularly marked impact on both baseline disease burden and intervention effectiveness.

We used empirical data to parameterise our model, but a noteworthy limitation is that these results may not be generalizable to other settings. However, we are not aware of any past work that has used empiric data to inform model parameters on the proportions of infectious TB and super-spreaders at all. As with any model-based analyses, our study has limitations that arise from its assumptions. In addition to those introduced above, our model is not intended to represent a specific setting, and as such does not incorporate stratification by age, HIV or multidrug-resistant TB. The other limitation is that we used a very simplified implementation of an active case finding intervention with a theoretical point-of-care diagnostic tool, without considering all the complexities in TB diagnosis and treatment implementation.

## Conclusions

The approaches we used to inform model parameters related to infectiousness heterogeneity from the observed distribution of secondary infections can be useful for future modelling studies of TB, in particular, and other infectious diseases, in general. The principles of implementation could be used in future TB modelling studies that represent specific epidemiological settings, especially when TB contact investigation data are available to estimate the proportion of super-spreaders. In the usage of advanced point-of-care technologies, targeting active case-finding interventions to super-spreaders is likely to be especially beneficial in low CDR settings.

## Acknowledgements

The authors would like to thank Monash University for providing the PhD scholarship to YAM.

## Availability of data and materials

The R-code used to analyse this model is publicly available.

## Authors’ contributions

YAM, RR and JMT conceived the study. YAM developed structure and ODE of the model, which was reviewed by JMT and RR. ESM, ACA, JMT conceived the strategy of the intervention in the model. YAM coded the model in R that RR and JMT reviewed and evaluated. YAM drafted the manuscript, and all authors provided input into revisions and approved the final draft for submission.

## Ethics approval and consent to participate

Not applicable

## Consent for publication

Not applicable

## Supplementary material

Supplementary Material. doc

